# The Integration of Pharmacophore-based 3D QSAR Modeling and Virtual Screening in Safety Profiling: a Case Study to Identify Antagonistic Activities against Adenosine Receptor, A2aR, using 1,897 Known Drugs

**DOI:** 10.1101/413385

**Authors:** Fan Fan, Dora Toledo Warshaviak, Hisham K. Hamadeh, Robert T. Dunn

**Author notes:** Currently in Comparative Biology and Safety Sciences, Amgen Inc., 360 Binney St., Cambridge, MA02142, USA. Currently in the Department of Molecular Engineering, Amgen Inc., 1 Amgen Center Dr. Thousand Oaks, CA 91320, USA. Currently in Technology Strategy and Innovation, Amgen Inc., 1 Amgen Center Dr. Thousand Oaks, CA 91320, USA. Currently in Charles River Laboratory, 334 South St, Shrewsbury, MA 01545.

## Abstract

Safety pharmacology screening against a wide range of unintended vital targets using *in vitro* assays is crucial to understand off-target interactions with drug candidates. With the increasing demand for *in vitro* assays, ligand-and structure-based virtual screening approaches have been evaluated for potential utilization in safety profiling. Although ligand based approaches have been actively applied in retrospective analysis or prospectively within well-defined chemical space during the early discovery stage (i.e., HTS screening and lead optimization), virtual screening is rarely implemented in later stage of drug discovery (i.e., safety). Here we present a case study to evaluate ligand-based 3D QSAR models built based on *in vitro* antagonistic activity data against adenosine receptor 2A (A_2a_R). The resulting models, obtained from 268 chemically diverse compounds, were used to test a set of 1,897 chemically distinct drugs, simulating the real-world challenge of safety screening when presented with novel chemistry and a limited training set. Due to the unique requirements of safety screening versus discovery screening, the limitations of 3D QSAR methods (i.e., chemotypes, dependence on large training set, and prone to false positives) are less critical than early discovery screen. We demonstrated that 3D QSAR modelling can be effectively applied in safety assessment prior to *in vitro* assays, even with chemotypes that are drastically different from training compounds. It is also worth noting that our model is able to adequately make the mechanistic distinction between agonists and antagonists, which is important to inform subsequent in vivo studies. Overall, we present an in-depth analysis of the appropriate utilization and interpretation of pharmacophore-based 3D QSAR models for safety screening.

## Introduction

Safety profiling against a wide range of molecular off-targets, prior to *in vivo* toxicity testing with animal models, has been widely implemented across the pharmaceutical industry[1-5]. Such a “bottom-up-approach” [6,7] reflects a continuous effort for a paradigm shift in early safety evaluations[8]. Besides preventing hazardous chemicals from entering animals, systematic screening is a necessary step to realize the vision of predicting human adverse events from mechanisms of action and the molecular targets involved. Safety profiling utilizes *in vitro* high throughput screens (HTS) against a broad array of unintended and vital targets. However as a safety screening panel typically includes a large number of targets, i.e., up to ~200[1,9], developing each liability target into a reliable HTS assay is resource demanding. As complementary approaches to help improve the utilization of *in vitro* HTS assays, tools such as ligand-and structure-based virtual screening have been evaluated. One advantage for *in silico* approaches is that it can be used to examine new compounds before they are synthesized, providing an attractive possibility for early hazard identification. If molecules with undesirable properties can be ruled out using *in silico* approaches, such as virtual screening, significant resources can be saved where only “prescreened” molecules are advanced to more costly *in vitro* screens.

For liability targets with little or no structural information, a ligand-based approach using quantitative structure activity relationship (QSAR) models may provide value[10-13]. QSAR is a machine learning process to develop meaningful correlations (model) between independent variables (e.g., structural features of compounds, molecular descriptors) and a dependent variable which is typically the value one wishes to predict [14]. The conceptual basis of such modeling is based on the hypotheses that compounds of similar structural features may exhibit similar biological activities[15]. A QSAR model is determined by factors such as activity data [16-18], molecule descriptors[16,19], and statistical algorithms[19,20]. Due to the advantages in throughput, cost saving(labor and reagents), turn-around-time, and the possibility to test compounds even before they are made, QSAR has been frequently used in various aspects of drug discovery such as lead optimization[14]. However, it has not been widely used in safety profiling, especially the 3D (i.e., pharmacophore) QSAR models, as most of commonly used QSAR models used in safety were built based on 2D features or molecular descriptors[21], such as the OECD QSAR toolbox[22], SEA[23], Toxmatch[24], ToxTreev[25], and DSSTox [26]. It is important to bear in mind the unique aspects for a safety profiling. In an efficacy screening (one target against many compounds), only the small amount of positives was considered. Quantitative determination of potency is crucial for lead optimization and ranking compounds. The negatives were of less value. Whereas in a safety profiling (often one compound against many targets), every data point counts including all negatives. In fact a negative result against a liability target for a drug candidate would be regarded as “good news”. As such, a false negative (contributing to sensitivity) result would be of greater concern in the safety space in comparison to a false positive (contributing to specificity), because it would mean advancing a potentially hazardous compound into further development. Quantitative value of potency is of less importance than efficacy screening. Due to these unique features and mindset, questions regarding QSAR applications remain in data interpretation as well as how to best incorporate these tools [27].

We present here a case study to evaluate the utilization of 3D QSAR modeling as a part of integrated approach to support safety profiling. Adenosine receptor 2a (A_2a_R) is one of the four class A GPCRs that regulate the activity of adenosine’s biological actions as a signaling molecule[28,29]. Due to its presence in both central nervous system and peripheral tissues[28,29], A_2a_R plays important roles in a wide range of biological processes such as locomotion, anxiety, memory, cognition, sleep regulation, angiogenesis, coronary blood flow, inflammation, and the anti-tumoral immunity [30-38]. Disruption of A_2a_R activities, consequentially, may result in undesired side effects in behavioral, vascular, respiratory, inflammatory, and central nervous systems. Indeed A_2a_R is a well-established liability target, as demonstrated in an industrial survey across four pharmaceutical companies [1]. Here, we developed QSAR models to predict compounds’ antagonistic activity against A_2a_R. It is important to note that the crystallographic structure of A_2a_R is known,^59^ in contrast to a large number of safety targets (e.g., ion channels and transporters). To make this study generalizable to those targets, however, we chose not to incorporate the structural data for A_2a_R in model building, but rather used it to provide additional insights to evaluate the performance of the ligand QSAR model. In our study, we collected 268 in house and external compounds with IC_50_ values against A_2a_R available, which were used to build the QSAR models. The majority of these compounds were obtained from early chemistry scaffolds and SAR. Hence, these compounds represented a diverse chemical space but not necessarily with ideal “drug-like features”. Bearing in mind that the goal is prospective utilization of QSAR in secondary pharmacology profiling, we tailored our study specifically within the setting of drug discovery. First, overtraining the model(s) was avoided. During drug development, it may not be practically possible to obtain many training compounds and assay results, hence the need to implement QSAR model. Therefore, we did not adhere the 4:1 or 10:1 ratio [39,40] for training and test sets. Second, as new chemotypes are constantly made in pharmaceutical development to drive SAR, a different external set of compounds were used to further challenge the QSAR models, as illustrated in **Fig 1**. This additional level of challenge came from 1,897 known drugs. Among these drugs, a subset of 75 known A_2a_R ligands was used as an external set. The 75 ligands in the subset are different in structure compared to the initial 268 training and test compounds. These 75 compounds were thoroughly tested to evaluate prospective utilization of the generated QSAR model(s) before applying them to screen the rest of ~1,800 drugs from the DrugBank[41]. These ~1,800 drugs further differ from the 268 compounds in chemical structure, which created a more realistic challenge. It is important to note that the focus of our study is the repurpose of existing QSAR tools in the realm of drug safety, rather than developing novel QSAR methodology. We demonstrate that, due to the unique requirements of safety screening, the well-known limitations of QSAR methods (i.e., chemotypes, dependence on large training set, and prone to false positives) are less critical than in early discovery screening. Overall, what we present is an in-depth case study for the utilization of *in silico* methods in early safety profiling.

## MATERIALS AND METHODS

### Materials

All compounds for *in vitro* assay validation were purchased from Sigma or Fisher Scientific where available. *Data set.* A total of 268 compounds were used as training (and test) set for pharmacophore-based 3D QSAR modeling. Among them, 87 compounds were downloaded from ChEMBL [42], and were then tested either by functional Ca^2+^ or cAMP assays. An additional 13 compounds were obtained from literature search and Guide to Pharmacology[43]. Another 168 internal compounds were selected based on our historical in-house Ca^2+^ flux assays. Chemical clustering analysis for these 268 compounds was performed using Schrödinger Canvas [44]. The pIC_50_ values, i.e., the negative logarithm values of IC_50_ were also calculated. Activity threshold of pIC_50_ = 5.0 was applied to set active from inactive compounds.

To validate the generated QSAR models, the SMILES codes and molecular descriptors of 1,806 approved and 179 withdrawn drugs were downloaded from the DrugBank [41]. After removing duplicates and applying the cutoff of 1,000 Da for molecular weight, 1,832 marketed drugs were obtained. An additional 65 known A_2a_R ligands, containing both agonists and antagonists, were downloaded from The Guide to Pharmacology as enrichment. These 65 compounds, along with 10 additional A_2a_R antagonists among the 1,832 Drugbank compounds, composed of a subset of 75 A_2a_R ligands, which were used to evaluate the performance of QSAR models. Collectively, 1,897 compounds were used as an external set.

## Methods

### Similarity comparison of chemical features between two sets of compounds

The radial binary fingerprints of chemicals were generated in Schrödinger Canvas (version 2.4), using the default settings according to the user manual. The subsequent comparison between sets were also carried out in Canvas using the Tanimoto similarity metrics, the resulting heat map of visualization was also generated.

### Chemical structure preparation

The 2D structures of all compounds in training, test, and external sets were converted to 3D using LigPrep (version 10.2, Schrödinger, LLC), using the default settings according to the user manual, where hydrogens were added, salts were removed, stereoisomers were generated, and the most probable ionization states were calculated at pH value of 7.0 ± 2.0 using the Epik module [45,46]. During ligand preparation, specified chirality was retained. As the conformations of the given compounds were unknown when bound to target, a series of 3D conformers (≤10 per rotatable bond, and ≤100 per ligand) were generated. The redundant conformers were eliminated using RMSD cut off value of 1.0 Å. The subsequent energy minimization of each structure was carried out using OPLS3 force field [47], and was filtered through a relative energy window of 10.0 kcal/mol to exclude high energy structures.

### Creating pharmacophore based models

The molecules were classified as actives and inactives by setting an activity threshold in Phase (actives: pIC_50_ ≥ 5.0) and inactives: pIC_50_ < 5.0). Each energy minimized ligand structure is described by a set of points (i.e., pharmacophore sites) in 3D space, representing various chemical features contributing to non-covalent binding between the ligand and the target of interest. These pharmacophore sites were characterized by type, position as well as directionality. Phase has 6 built-in pharmacophore types: hydrogen bond acceptor (A), hydrogen bond donor (D), hydrophobe (H), negative ionizable (N), positive ionizable (P), and aromatic ring (R). Pharmacophore features that are common to most actives (e.g., ≥ 50%) were identified to perceive pharmacophore hypotheses.

Such generated pharmacophore hypotheses were then scored based on the superimposition of the site points, vector alignment and volume overlap [48]. Scoring was obtained first with all active compounds, and then inactive compounds. The hypotheses that matched the inactive ligands were penalized as described in details by Dixon *et al* [48]. Default values were used for weights (w) of actives and inactives.

### Pharmacophore-based 3D QSAR modeling

3D QSAR models were generated using atom-based PLS (partial least square) regression method. The default value of PLS of 3 was applied. For each of the top scored pharmacophore hypothesis, a QSAR model was built using training compounds that matched the pharmacophore on at least 3 sites and yielded best alignments [48]. Specifically, to generate a QSAR model, a rectangular grid was defined to include the space occupied by the aligned training set actives. The grid was divided into uniformly sized cubes of 1 Å^3^. The cube was deemed as occupied if the center of a pharmacophore site was within the radius of the corresponding sphere. Based on the differences in the occupancy of cubes and the different types of sites that reside in these cubes, a compound may therefore be represented by a string of zeros and ones. This resulted in binary values as 3D descriptors. QSAR models were created by using partial least square regression (PLS) to the pool of binary valued variables [48].

The generated QSAR models were examined using the test set compounds. By comparison of the predicted and experimentally determined pIC_50_ values, the statistical parameters R^2^ (correlation coefficient), SD (standard deviation of regression) and Root Mean Square Deviation (RMSD) were calculated to evaluate the overall significance of the model. The best performed model was selected for the subsequent virtual screen.

### Pharmacophore-based 3D QSAR virtual screening

The 1,897 known drugs were energy minimized and conformations were generated to form the 3D database (library) in Phase. The pharmacophore hypotheses of the best 3D QSAR models were used to screen against this library for compounds that match such pharmacophore features. The pIC_50_ values of the hit compounds were then predicted using the 3D QSAR model.

***In vitro* Assays**.

Competition binding assay using radioactive ligands and functional assay monitoring cAMP were carried out as a paid service provided by CEREP (Poitiers, France). The functional assays monitoring Ca^2+^ flux were carried out as a paid service provided by DiscoveRx (Carlsbad, CA). The competition binding assays were carried out at a fixed compound concentration of 10 μM. The cAMP and Ca^2+^ assays were carried out in concentration-response mode, at 10, 3.165, 1.001, 0.317, 0.100,0.003, 0.001, and 0 μM.

## RESULTS

### Data preparation and chemical clustering

The 268 compounds, from public sources and in-house, were divided into training and test sets via 3 approaches as illustrated in **Fig 1**, as QSAR outcome might be affected by how training and test sets were separated [40,49]. In the first approach, a randomized separation was used, resulting in 141 training compounds, and 127 test compounds. Cheminformatics analysis revealed that the training and test sets contained 26 and 37 chemical clusters, respectively, among which only 8 clusters were shared between the 2 sets. A second method to create a training set covering more chemical clusters was also applied. As the 268 compounds contained 55 chemical clusters, the centroids of the 55 chemical clusters (among which only 8 are actives), and randomly selected 27 actives, were grouped into the training set. The remaining 185 compounds were used as the test set. In the third approach, all 268 compounds were used as one training set to create common pharmacophore hypotheses. To make a distinction, the training and test sets obtained from random separation were referred to as training 1 and test 1. The training and test sets from the clustering method were referred to as training 2 and test 2. The training set from the third approach was referred to as training 3. The obtained hypotheses from these training sets were subsequently used to build pharmacophore-based 3D QSAR models.

**Fig 1.**
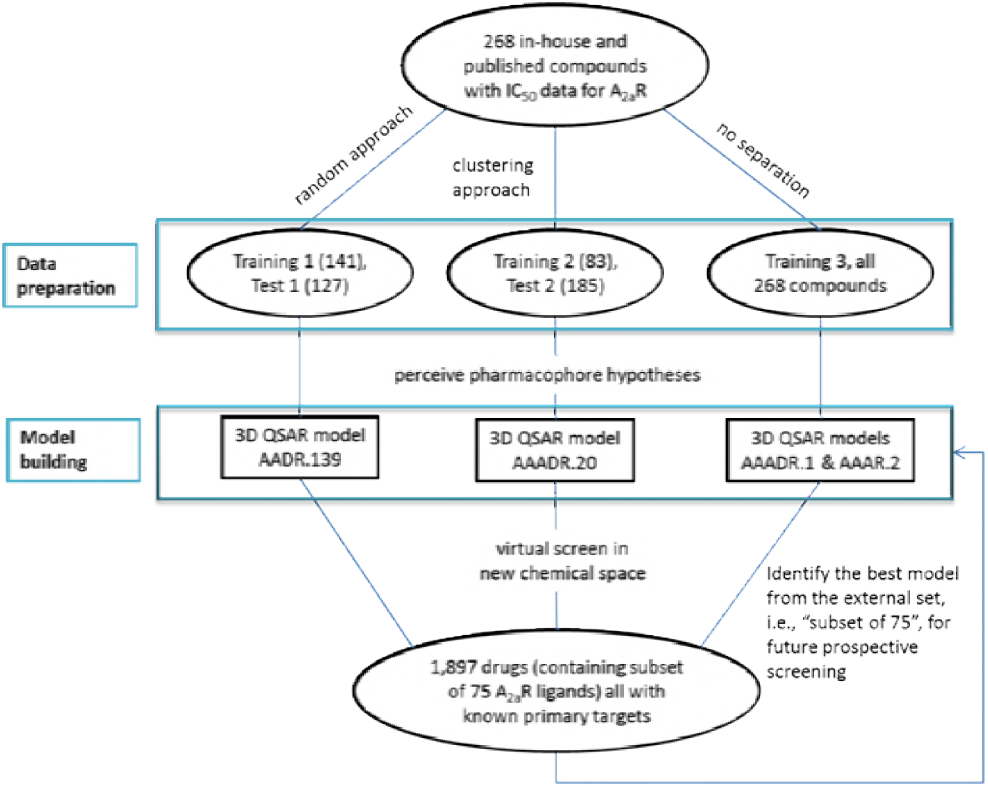
Workflow illustration for pharmacophore-based 3D QSAR modeling and virtual screening to identify compounds with antagonistic activities against A_2a_R.

The similarity analysis between training and test sets was carried out using Canvas [44]. Training 1 and test 1 exhibited some difference in their chemical features, as revealed by the similarity index, ranging from 0.01 to 0.65 (**Fig 2A**). As shown in the heat map in **Fig 2B**, the similarity index between the training 2 and test set 2 compounds ranged from 0.01 to 0.67.

The 1,897 compounds bore less similarity with the 268 compounds, which were represented by the 55 centroids. The similarities with the 55 representatives ranged from 0.01 to 0.37, and 0.01 to 0.13, with the subset of 75 A2aR ligands (**Fig 2C**) and the 1,832 compounds (**Fig 2D**), respectively. This is perhaps not entirely surprising, as the 268 compounds represented chemical space at the discovery stage, whereas the 1,897 molecules represented the true drug space. Due to low structural similarities, the screening of 1,897 drugs using pharmacophore-based 3D models generated from early stage chemicals presented a “real world” case scenario in Drug Discovery.

**Fig 2.**
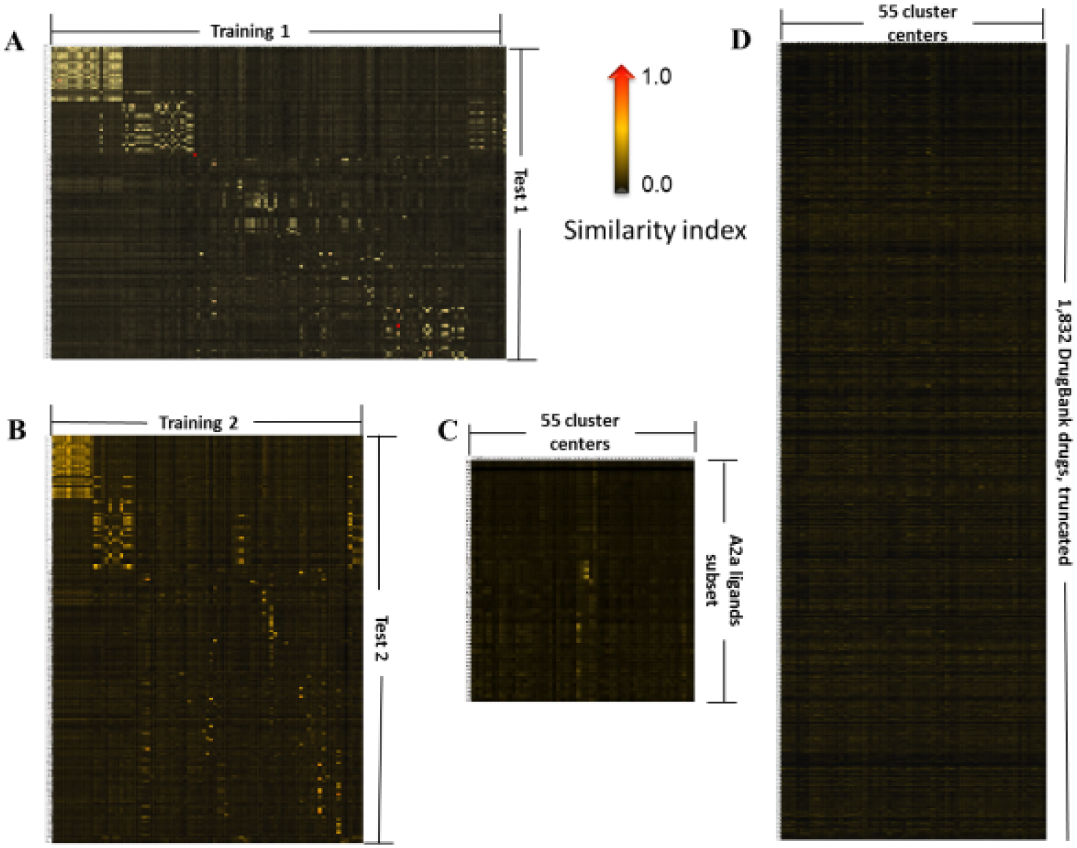
The heat map demonstration for binary fingerprint similarities between training 1 and test 1 (**A**), training 2 and test 2 (**B**), the 55 representations of the 268 compounds and the subset of 75 A_2a_R ligands (**C**), as well as the 55 representations and the 1,832 DrugBank drugs (**D**). The heat map of 1,832 drugs was truncated due to space limitation. The heat maps were generated using Schrodinger Canvas, as described in details in Materials and Methods. The lowest similarity (0.0) was shown in black, whereas the highest similarity (1.0) was shown in red. See supplementary data for a zoomed in version for each panel.

**Pharmacophore modeling**.

The training set compounds were divided into actives (pIC_50_ ≥ 5.0), and inactives (pIC_50_ < 5.0), consistent with our in house *in vitro* profiling practice. Training sets 1, 2, 3 contained 54, 35 and 97 actives, respectively. Various combinations of common pharmacophores were identified. From training set 1, 6 four-pharmacophore-site variants were generated to match ≥ 40 out of 53 actives. From training 2, a total of 7 five-pharmacophore-site variants were generated to match ≥ 21 of the 35 actives. Only 3 five-pharmacophore-site variants were generated to match ≥ 55 out of 97 actives in training 3. The possibility of four-pharmacophore-site variants was also explored, from which 8 variants were produced to match ≥ 63 out of 97 actives. The variants and the possible resulting hypotheses were summarized in **Table 1**.

Upon completion of scoring for all the hypotheses listed in **Table 1**, 46 four-site hypotheses survived from training 1, 9 five-site survived from training 2. In training set 3, 4 five-site hypotheses survived and 19 four-site hypotheses survived. The top survived hypotheses (~10%-~20%)were used to build 3D QSAR models.

**Table 1.**
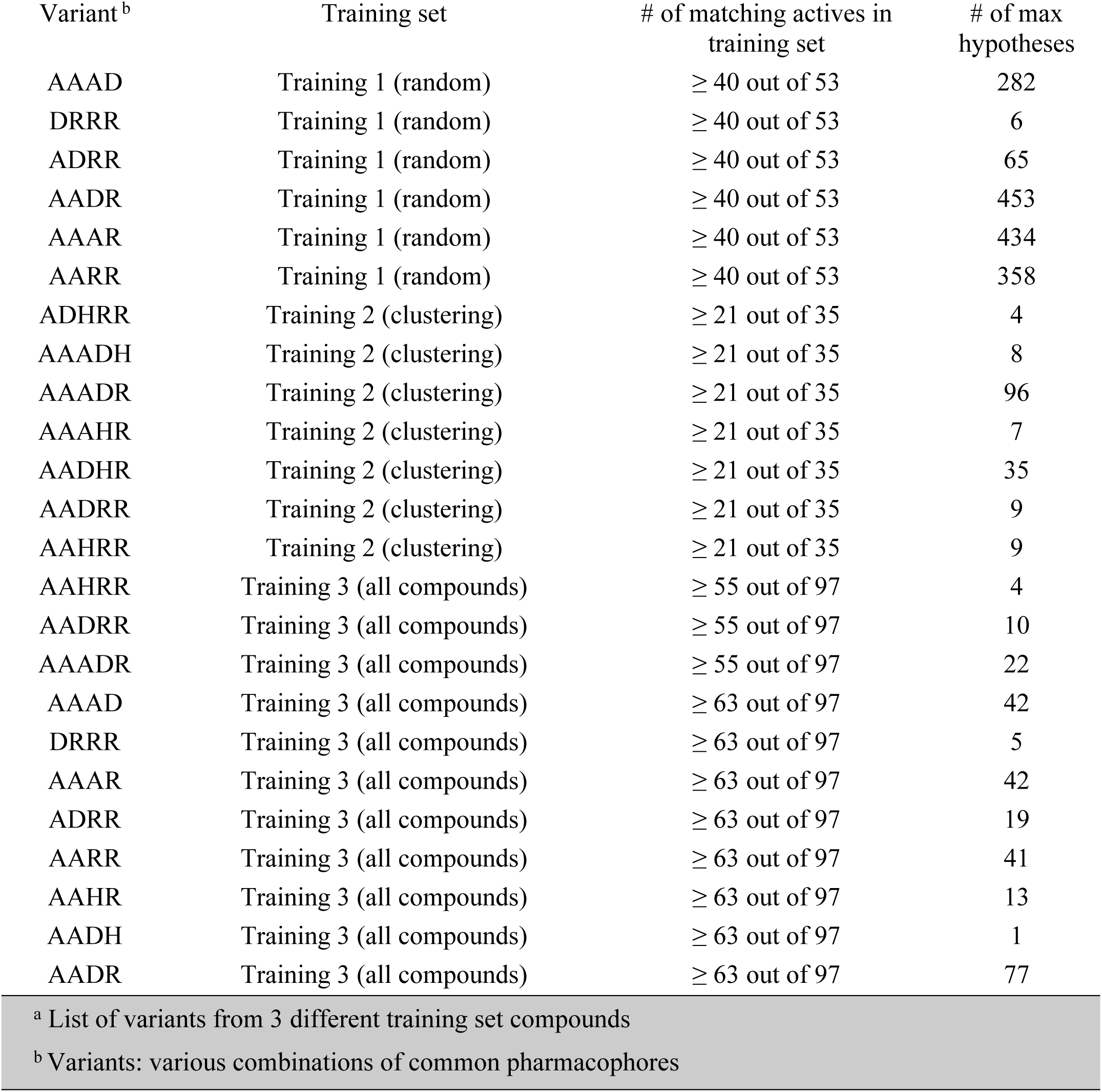
Pharmacophore hypotheses identified by Phase.

**Generation and test of the Pharmacophore-based 3D QSAR model**.

Four and three pharmacophore models were generated from the survived hypotheses, for training 1 and 2, respectively. Evaluation of these models was performed by predicting activities for test 1 and 2 sets of compounds. As the number of PLS factors increased, the statistical significance and predictive ability of the model was also incrementally increased. Therefore, PLS factor of 3 were used for the models. The statistical results were summarized in **Table 2**. It was found that AADR.139 and AAADR.20 yielded the best statistics for test 1 and test 2, respectively. The large F value and the small p value indicated a statistically significant regression model and high degree of confidence. The small value of SD and RMSE suggested satisfactory results from the test set. The q^2^ value was indicative of the capability to predict activities in the test set. The performance of predicting activities of the test set could also be seen from the correlation between predicted and experimentally determined pIC_50_ values as shown in **Fig 3**. Both AADR.139 and AAADR.20 were moved forward to generate 3D QSAR models. For model AADR.139, sensitivity and specificity were observed to be 82% and 94% against test set 1; for model AAADR.20, the sensitivity and specificity against test set 2 reached 96% and 94%, respectively.

**Table 2.**
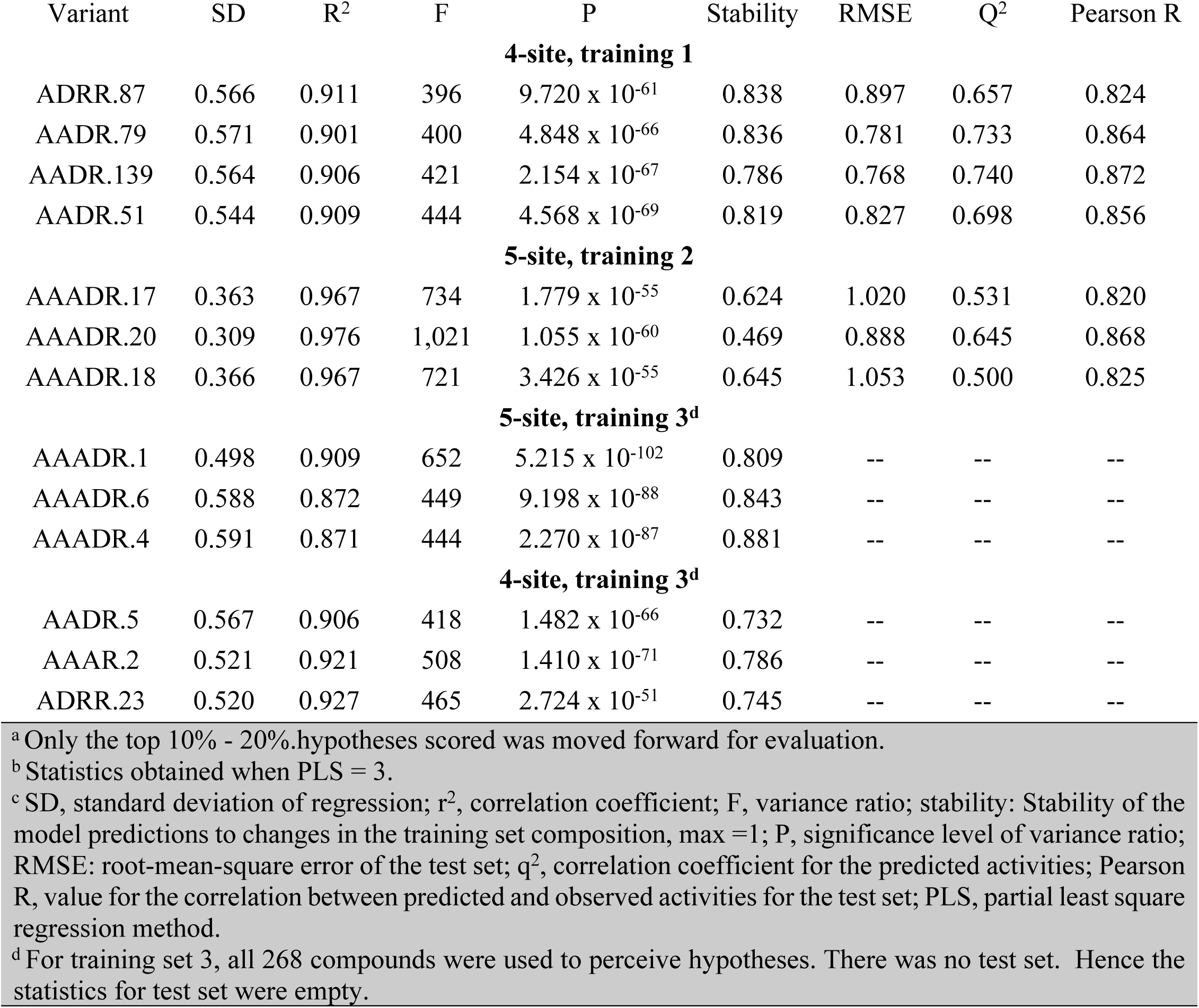
The statistical data of pharmacophore-based 3D QSAR using Phasea,b,c.

**Fig 3.**
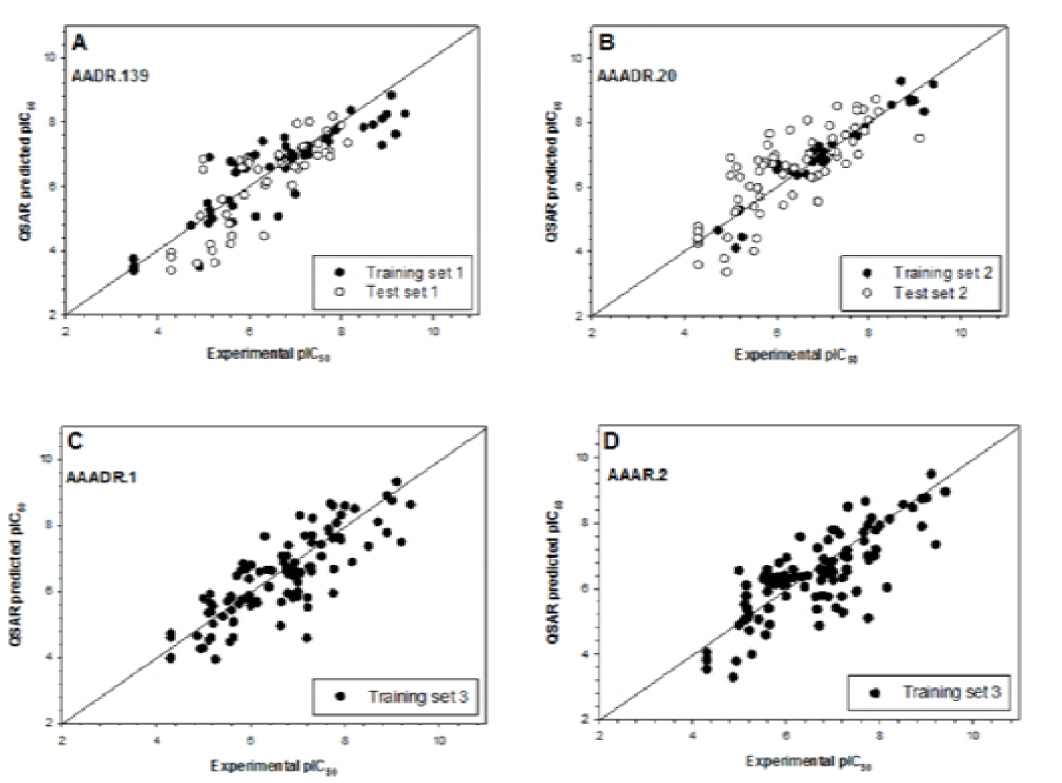
The performance of the 4 models on predicting actives of the test set compounds. **A**, model AADR.139 generated from training set 1. **B**, model AAADR.20 generated from training set 2. **C**, model AAADR.1 generated from training set 3. **D**, model AAAR.2 generated from training set 3. In the cases of models AAADR.1 and AAAR.2, there were no test set compounds as all 268 compounds were used as training, as described in details in Materials and Methods.

From training set 3, three 3D pharmacophore models were generated for five-site and four-site hypotheses. Unlike the previous 2 training sets, there is no test set for training 3. Nonetheless, AAADR.1 and AAAR.2 were the best five-site, and four-site models, respectively, as shown by the statistical parameters that are unrelated to the test set. It is worth noting that AAAR.2 and AADR.5 shared the same reference compound (N-isopropyl-2-((pyridin-3-ylmethyl)amino)thieno[3,2-d]pyrimidine-4-carboxamide), which is the compound that matches the hypothesis with the highest score. N-Isopropyl-2-((pyridin-3-ylmethyl)amino)thieno[3,2-d]pyrimidine-4-carboxamide contained all five pharmocophore sites, 3 hydrogen bond acceptors, 1 hydrogen bond donor and 1 aromatic residue. It is therefore interesting to perform the subsequent virtual screen with both five-site and four-site pharmacophore models generated from the training set 3. The reference compound for each model was shown in **Fig 4**.

**Fig 4.**
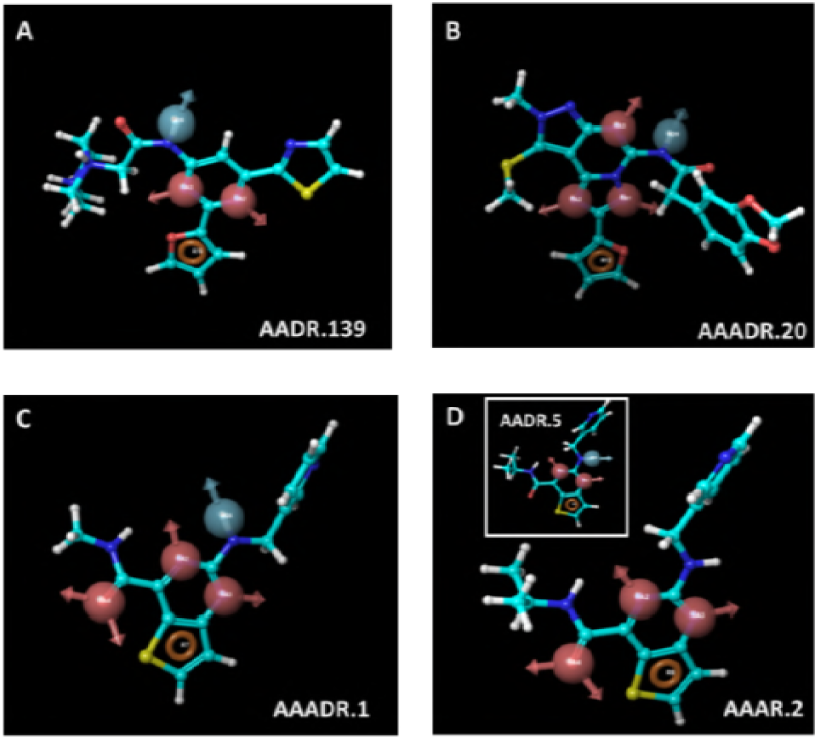
The reference ligands used for model AADR.139 (**A**), AAADR.20 (**B**), AAADR.1 (**C**), and AAAR.2 (**D**). The insert of Fig 4D shows that model AADR.5 shared the same reference compound (*N*-isopropyl-2-((pyridin-3-ylmethyl)amino)thieno[3,2-d]pyrimidine-4-carboxamide) as model AAAR. 2.

Hence, *N*-isopropyl-2-((pyridin-3-ylmethyl)amino)thieno[3,2-d]pyrimidine-4-carboxamide contained all five pharmocophore sites in model AAADR.1, as described in details in Results. Hydrogen bond acceptor was shown in magenta vector, hydrogen donor was shown in light blue vector, and aromatic residues were shown in brown ring.

## Virtual screening of 1,897 drugs

The 4 best models, built from 3 training sets containing 83 to 268 compounds, were used to perform virtual screening against 1,897 drugs. The 1,833 known drugs from DrugBank contained only 2 known A_2a_R agonists and 8 antagonists. Therefore, an additional 67 A_2a_R ligands (either drugs or drug candidates) were downloaded from Guide to Pharmacology. After removing the 2 duplicated molecules (A_2a_R agonists: Regadenoson and Adenosine), a 1,897-compound set was obtained. The 1,897-compound set contained 29 A_2a_R agonists and 46 antagonists. These 75 known A_2a_R ligands were used to evaluate the performance of the 4 models. With 6 CPUs, the screening process against 1,897 compounds with various conformers was completed within 3 minutes. The compounds which yielded predicted pIC_50_ values equal or larger than 5.0 were labeled as hits. The number of hits and hit rates using each model was summarized in **Table 3**.

**Table 3.** Virtual screen hits and hit rates obtained from 4 different pharmacophore-based 3D QSAR models.

| Model | AADR.139 | AAADR.20 | AAADR.1 | AAAR.2 |
| --- | --- | --- | --- | --- |
| Complete set, 1,897 compounds |  |  |  |  |
| # of Hits | 115 | 77 | 83 | 168 |
| Hit rate, % | 6.1 | 4.1 | 4.4 | 8.9 |
| Subset, 75 A<sub>2a</sub>R ligands (29 agonists & 46 antagonists)<sup>a, b</sup> |  |  |  |  |
| Sensitivity, % | 28 | 28 | 13 | 78 |
| Specificity, % | 89 | 78 | 45 | 74 |
| False positive rate, % | 11 | 22 | 55 | 26 |
| False negative rate, % | 72 | 72 | 87 | 22 |

<sup>a</sup> Performance statistics were calculated based on the assumption that agonists should behave as negatives, i.e., yielding pIC<sub>50</sub> < 5.0 when being tested via functional antagonist assay.
<sup>b</sup> Sensitivity = TP/(TP+FN), specificity = TN/(TN+FP), false positive rate = FP/(FP + TN), false negative rate = FN/(TP+FN), where TP is true positive, TN is true negative, FP is false positive, FN is false negative.

As demonstrated from the subset containing 75 A_2a_R ligands, model AAAR.2 yielded > 70% sensitivity and specificity, a significantly better performance in comparison to the other 3 models, despite the assay statistics (sensitivity and specificity) may be underestimated. All subsequent discussions will be focused on AAAR.2. Some active antagonists, such as MRS1532, MRS1191, MRS1088, dyphyllines, and pentoxiphylline etc., were weak antagonists with pIC_50_ values around 5.0. These compounds were predicted to be inactives (predicted pIC_50_ < 5.0) by the model, causing them to be categorized as “false negatives” and in turn underestimating the sensitivity. On the other hand, the agonists were viewed as negatives in the antagonistic A_2a_R QSAR model. In our screening, 7 agonists had predicted antagonistic pIC_50_ ≥ 5.0. These 7 agonists were deemed as false positives in our analysis. However, it is not unusual for agonists to demonstrate antagonistic activities [50], which was indeed observed in the *in vitro* assays with adenosine and regadenoson (*vida infra*). Accordingly, the false positives might be overestimated, which may in turn result in underestimated specificity. It is also important to note that AAAR.2 was the only model that successfully identified 8 out of 9 drugs in the theophylline family. The only family member failed to be identified was fenethylline, as AAAR.2 predicted doxofylline and pentoxifylline to have activity below the pIC_50_ cutoff of 5.0 (pIC_50_ predicted to be 4.2 and 3.7 respectively).

Given the structural difference between the subset and the 268 training (and testing) compounds, AAAR.2 was determined to be the most relevant pharmacophore-based 3D QSAR model based on its performance. The 3-dimensional aspects of the QSAR model AAAR.2 were further examined to help gain an understanding on how the structures of the ligands contribute to the A_2a_R antagonistic activities. The intersite distance between pharmacophores is shown in **Fig 5**. The 4 pharmacophores, A2, A3, A4 and R8, formed a diamond shape, with the longest distance (5.1 Å) occurring between A3 and A4. The positive and negative coefficients that contribute to the increase or decrease in antagonistic activity against A_2a_R could be visualized by pictorial representations (**Fig 6**). The blue cubes indicated the favorable regions for a given feature, whereas the red cubes indicated unfavorable regions. **Fig 6A** highlighted the favorable and unfavorable regions for the presence of hydrogen bond donor. Similarly, the favorable and unfavorable regions for hydrophobic groups and electron withdrawing groups were shown in **Figs 6B** and **6C**, respectively. The combined, overall effects were shown in **Fig 6D**. The reference ligand, N-isopropyl-2-((pyridin-3-ylmethyl)amino)thieno[3,2-d]pyrimidine-4-carboxamide, was primarily covered in favorable regions, especially the core area composed of the 4 pharmacophores despite some unfavorable regions around the edges. The only major exception was around the methylpyridine moiety, whose hydrophobicity was disfavored for the antagonistic activity. The reference ligand had a moderate activity (pIC_50_ = 5.6), within a concentration range commonly observed in Safety profiling. Such visualization is also useful to examine some more active and inactive ligands to identify structural features that may be unfavorable, as detailed in the supplementary data.

**Fig 5.**
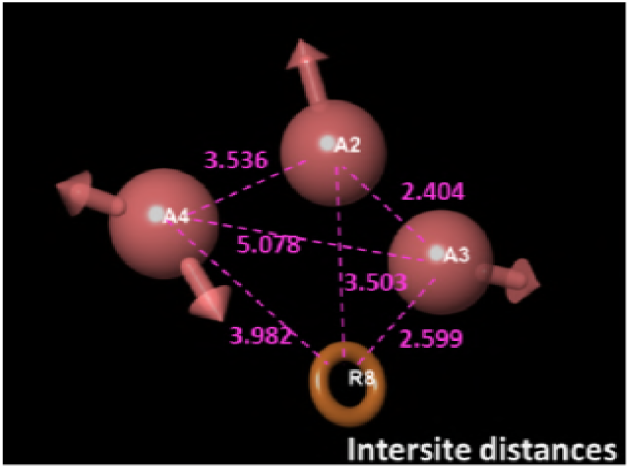
The inter-site distances between model AAAR.2. Distances are in the unit of Å.

**Fig 6.**
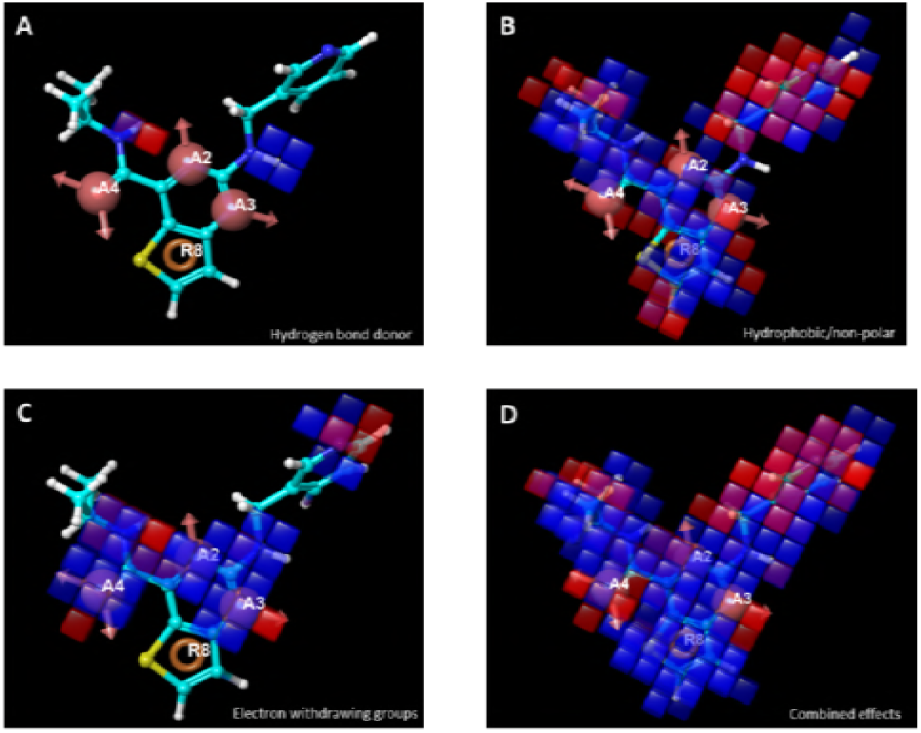
Pictorial representations of the positive (cobalt) and negative (red) coefficients that contribute to A_2a_R antagonist activities, from hydrogen bond donor (**A**), hydrophobicity (**B**), electron withdrawing groups (**C**), and the combined effects (**D**).

Further validation of virtual screen results using *in vitro* assays. A total of 56 randomly selected compounds from virtual screening were subjected to cross validation using in vitro assays using three different assay formats. The 56 compounds includes compounds from 3 categories: 16 with predicted pIC_50_ ≥ 5.0 (virtual screen actives), 17 yielded predicted pIC_50_ ranging from 4.0 to 5.0, and 23 compounds either yielded predicted pIC_50_ < 4.0 or were not even picked up by the screening (virtual screen inactives). As shown in **Table 4**, only 5 out of the 16 virtual screen actives, i.e., amiloride, theophylline, doxorubicin, S-adenosylmethionine, and pranlukast, were confirmed by at least 1 type of *in vitro* assay. Among the virtual screening negatives, both A_2a_R agonists, adenosine and regadenoson were picked up by AAAR.2, although the predicted pIC_50_ were 4.7 and 4.9, respectively. All other negatives were confirmed as negatives by in vitro assays.

**Table 4.** *In vitro* assay results

| Molecule | IC<sub>50</sub>-<br>predicted | IC<sub>50</sub>,<br>cAMP,<br>M | IC<sub>50</sub>,<br>Ca<sup>2+</sup>, M | %inhibition,<br>binding* | Primary target(s) and/or<br>function(s) |
| --- | --- | --- | --- | --- | --- |
| Amiloride | 4.0E-07 | N.E. | N.E. | 53 | Amiloride-sensitive Na<sup>+</sup> channel<br>subunit $\alpha$ |
| Theophylline | 5.0E-07 | N.E. | N.E. | 68 | A<sub>2a</sub>R antagonist |
| Triamterene | 6.3E-07 | N.E. | N.E. | N.E. | Amiloride-sensitive Na<sup>+</sup> channel |
| S-Adenosyl-<br>methionine | 7.9E-07 | N.E. | N.E. | 74 | common cosubstrate for<br>methyl transferase and so on |
| Pranlukast | 1.0E-06 | N.E. | 8.3E-07 | N.E. | cysteinyl leukotriene receptor 1<br>antagonist |
| Valganciclovir | 1.3E-06 | N.E. | N.E. | N.E. | a prodrug for ganciclovir, DNA,<br>transporters |
| Bortezomib | 1.6E-06 | N.E. | N.E. | N.E. | proteasome inhibitor |
| Ribavirin | 1.6E-06 | N.E. | N.E. | N.E. | adenosine kinase, Inosine-5'-<br>monophosphate dehydrogenase<br>1 inhibitor |
| Ticagrelor | 2.5E-06 | N.E. | N.E. | N.E. | P2Y, platelet aggregation<br>inhibitor |
| Famotidine | 4.0E-06 | N.E. | N.E. | N.E. | H2 receptor antagonist |
| Valaciclovir | 4.0E-06 | N.E. | N.E. | N.E. | thymine kinase inducer, DNA<br>polymerase inhibitor |
| Cefdinir | 6.3E-06 | N.E. | N.E. | N.E. | β-lactam antibiotic |
| Salsalate | 6.3E-06 | N.E. | N.E. | N.E. | Prostaglandin G/H synthase<br>1&2 |
| Pyridoxal | 6.3E-06 | N.E. | N.E. | N.E. | Pyridoxal kinase, precursor to<br>pyridoxal phosphate |
| Doxorubicin | 7.9E-06 | N.E. | 2.5E-06 | N.E. | DNA intercalator, DNA topoisomerase inhibitor |
| Bopindolol | 7.9E-06 | N.E. | N.E. | N.E. | $\beta$ blocker |
| Cefixime | 1.3E-05 | N.E. | N.E. | N.E. | β-lactam antibiotic |
| Adenosine monophosphate | 1.3E-05 | N.E. | 1.9E-07 | 65 | A<sub>2a</sub>R agonist |
| Regadenoson | 1.6E-05 | N.E. | 4.9E-08 | 97 | A<sub>2a</sub>R agonist |
| Sofosbuvir | 2.0E-05 | N.E. | N.E. | N.E. | prodrug nucleotide analog |
| Capecitabine | 2.0E-05 | N.E. | N.E. | N.E. | Prodrug of 5-FU, Thymidylate synthase inhibitor |
| Milrinone | 2.0E-05 | N.E. | N.E. | N.E. | cAMP phosphodiesterase inhibitor |
| Nebivolol | 2.0E-05 | N.E. | N.E. | N.E. | β1 receptor antagonist |
| Reboxetine | 2.0E-05 | N.E. | N.E. | N.E. | Na<sup>+</sup>-dependent noradrenaline transporter inhibitor |
| Propafenone | 3.2E-05 | N.E. | N.E. | N.E. | Na<sup>+</sup>, K<sup>+</sup> channels blocker |
| Felbamate | 3.2E-05 | N.E. | N.E. | N.E. | NMDA receptors antagonist |
| Flucloxacillin | 3.2E-05 | N.E. | N.E. | N.E. | β-lactam antibiotic |
| Sulpiride | 3.2E-05 | N.E. | N.E. | N.E. | D2 antagonist |
| Bosentan | 4.0E-05 | N.E. | N.E. | N.E. | endothelin receptor antagonist |
| Ipratropium bromide | 4.0E-05 | N.E. | N.E. | N.E. | Muscarinic receptor antagonist |
| Gliquidone | 5.0E-05 | N.E. | N.E. | N.E. | ATP-sensitive K<sup>+</sup>-channel inhibitor |
| Metoprolol | 6.3E-05 | N.E. | N.E. | N.E. | β1 blocker |
| Glimepiride | 7.9E-05 | N.E. | N.E. | N.E. | ATP-sensitive K<sup>+</sup>-channel receptor inhibitor |
| Midodrine | 1.0E-04 | N.E. | N.E. | N.E. | alpha-adrenergic receptor agonist |
| Norepinephrine | 1.0E-04 | N.E. | N.E. | N.E. | alpha-adrenergic receptor agonist |
| Isradipine | 1.0E-04 | N.E. | N.E. | N.E. | calcium channel blockers |
| Pentoxifylline | 2.0E-04 | N.E. | N.E. | N.E. | Phosphodiesterase inhibitor, adenosine receptor antagonist |
| Verapamil | 3.2E-04 | N.E. | N.E. | N.E. | L type Ca<sup>2+</sup> channel inhibitor |
| Diltiazem | 7.9E-03 | N.E. | N.E. | N.E. | L type Ca<sup>2+</sup> channel inhibitor |
| Cefadroxil | non hit | N.E. | N.E. | N.E. | β-lactam antibiotic |
| Nelfinavir | non hit | N.E. | N.E. | N.E. | HIV-1 protease inhibitor |
| Cephalexin | non hit | N.E. | N.E. | N.E. | β-lactam antibiotic |
| Rosiglitazone | non hit | N.E. | N.E. | N.E. | PPARγ agonist |
| Cefoxitin | non hit | N.E. | N.E. | N.E. | β-lactam antibiotic,<br>carboxypeptidase inhibitor |
| Etravirine | non hit | N.E. | N.E. | N.E. | Non-Nucleoside Reverse<br>Transcriptase Inhibitor |
| Pirlindole | non hit | N.E. | N.E. | N.E. | Non-Nucleoside Reverse<br>Transcriptase Inhibitor |
| Desogestrel | non hit | N.E. | N.E. | N.E. | synthetic progestational<br>hormone |
| Pheniramine | non hit | N.E. | N.E. | N.E. | H1 antagonist |
| Gabapentin | non hit | N.E. | N.E. | N.E. | Voltage-gated Ca<sup>2+</sup> channel<br>inhibitor |
| Ticlopidine | non hit | N.E. | N.E. | N.E. | P2Y antagonist |
| Mesalazine | non hit | N.E. | N.E. | N.E. | Prostaglandin G/H synthase<br>1&2 inhibitor |
| Flumethasone | non hit | N.E. | N.E. | N.E. | GR agonist |
| Cabergoline | non hit | N.E. | N.E. | N.E. | dopamine agonist, prolactin<br>inhibitor |
| Lamotrigine | non hit | N.E. | N.E. | N.E. | Voltage-gated Na<sup>+</sup> channel<br>inhibitor |
| Nitisinone | non hit | N.E. | N.E. | N.E. | 4-Hydroxyphenylpyruvate<br>dioxygenase inhibitor |
| Ertapenem | non hit | N.E. | N.E. | N.E. | β-lactam antibiotic |

\*% of inhibition = $100 - \left( \left( \frac{\text{Measured specific binding}}{\text{control specific binding}} * 100 \right) \right)$ . Compound binding was calculated as a % inhibition of the binding of a radioactively labeled ligand specific for each target.
\*\* N.E., no effects. Results showing an inhibition or stimulation lower than 50% are considered to represent insignificant effects of the test compounds.

The compounds were determined as “active” when confirmed by at least 1 type of in vitro assay. Identifying actives using 3 different assays was to reduce any “omissions” (false negatives) caused by artifacts with any one form of particular assay. The sensitivity and specificity of AAAR.2 for predicting the random 56 drugs are summarized in **Table 5**.

**Table 5.** Performance of prediction and chemical similarities^a,b^

| Model | Training 1<br>vs Test 1 | Training 2<br>vs Test 2 | Training 3 vs 75<br>A2aR ligands | Training 3 vs 56<br>drugs b |
| --- | --- | --- | --- | --- |
| # of clusters in training | 26 | 55 | 55 | 55 |
| # of actives in training | 53 | 35 | 97 | 97 |
| # of inactives in training | 88 | 48 | 171 | 171 |
| Max similarity to test set <sup>c</sup> | 0.65<sup>c</sup> | 0.67 | 0.37 | 0.13 |
| Sensitivity, % | 82 | 96 | 78 | 72 |
| Specificity, % | 94 | 94 | 74 | 77 |

<sup>a</sup> Sensitivity = TP/(TP+FN), specificity = TN/(TN+FP), false positive rate = FP/(FP + TN), false negative rate = FN/(TP+FN), where TP is true positive, TN is true negative, FP is false positive, FN is false negative.
<sup>b</sup> Based on in vitro assay results, TP =5, TN =38, FP =11, FN = 2.
<sup>c</sup> Similarities calculated using radial binary fingerprints. The 268 training and test compounds were represented by 55 centeroid structures from the 55 chemical clusters.

## DISCUSSION

Selectivity screening is an essential step to realize the vision of predicting the adverse events in human from molecular targets, and ultimately design away from these liability targets. With the increasing demand for *in vitro* assays as well as the expanding list of liability targets, tools such as ligand-and structure-based virtual screens have been evaluated to aid and optimize the profiling process in the realm of Predictive Safety. For targets whose structures are not available or for targets whose binding sites are flexible, ligand-based approach provides a powerful predictive tool, especially with carefully curated training and test compounds. *In silico* approaches in safety profiling are still at an early stage as questions remain in data interpretation as well as how to best incorporate these tools [27]. From the presented case study of screening antagonistic activity against A_2a_R, we evaluated how to best use and interpret pharmacophore-based 3D QSAR model in Safety.

### Data collection for model building

The success of modeling requires large and diverse training sets. Many researchers suggested a 10:1 or 4:1 ratio for the numbers of compounds in training and test sets [39,40] Although a significantly sized training set will help greatly in a QSAR exercise, in the reality of drug discovery particularly in safety, it is not always attainable to generate large quantity of *in vitro* data upfront. The notion of requiring *in vitro* data for large number of compounds indeed hampers the prospective utilization of QSAR models in the pharmaceutical industry. Besides, if a particular *in vitro* assay was readily available and new synthetic chemistry was quickly worked out, there would be practically no need to use *in silico* approaches. The most frequent question is: how many compounds are enough? Although it is not possible to put a fixed number, it would help to know how many compounds will be screened in the prospective utilization. Our study presented an extreme case using 268 training/test compounds to screen 1,897 compounds, demonstrating that the possibility of utilization QSAR even with smaller training/test set.

We suggested a couple of mitigation solutions to use QSAR when training and test sets are smaller than future screening task. One is to include data from public databases, such as ChEMBL[42] and Guide to Pharmacology [43]. Although these external compounds may represent different chemistry compared to an in-house produced collection, incorporation of these compounds enhanced chemical diversity. The second is to split training and test sets in various ways. As shown in our study, splitting training and test sets in various ways impacted the performance with external set (**Table 3**). Changing composition of the training and test set disturbed the basis of modeling, which in turn alters the outcomes [40,49]. Last, generating more than one hypothesis and model from each training set may be beneficial. Using the same training 3, various hypotheses yielded different statistics. For example, when changing AAADR.4 to AAADR.1, the compounds accounted for in the training set increased from 87% to 91%, as shown in the R^2^ values (Table 2). When changing from 5-site to 4-site hypotheses, the coverage further increased to 93%.

### The 4-site AAAR.2 is a relevant model to mechanistically predict new molecules for A_2a_R

Up to 7 pharmacophore sites can be defined in Schrödinger Phase [48,51]. Typically increasing numbers of pharmacophore site renders additional definitions for ligand features, which may better distinguish actives from inactives. While this might be true with more rigid binding pocket such as kinases [52], it was not the case for A_2a_R as demonstrated in our study. With training set 3, several 5-and 4-site models were obtained, all of which yielded good statistics as shown in **Table 3**. Despite the significantly improved *P* values in 5-site model, the 4-site model AAAR.2 yielded a significantly improved outcome when predicting the structurally different subset of 75 A_2a_R ligands. As revealed by our training set, log*P* values of the active compounds ranged from 0.2 to 7.2, suggesting ligands with a broad diversity are able to bind to this target. Indeed promiscuity is well known for target classes such as GPCR and nuclear hormone receptors [50,53,54]. Fewer pharmacophore sites may instead allow more freedom for the structurally “fluid” GPCRs. Therefore, it is important to test hypotheses composed of different number of pharmacophore sites, and evaluate the resulting models in the external set.

Model AAAR.2 was determined to be the most relevant pharmacophore model based on its performance against the 75 A_2a_R ligands and the 56 randomly selected drugs, both of which are structurally very different comparing to the 268 training compounds. AAAR.2 contained 4 pharmacophore features, 3 hydrogen bond acceptors and 1 aromatic ring. The emphasis for hydrogen bond acceptors can be seen from the 97 actives in training set 3, among which the number of hydrogen bond acceptor ranged from 3 to 8. In contrast, the presence of a hydrogen bond donor was not necessary for antagonistic activities, as 9 out of 97 actives contained no hydrogen bond donors. An in-depth survey for a database (SCOPE database[55]), containing proprietary compound optimization data, showed that average number of hydrogen bond acceptors for GPCR ligands increased from 3 to 4 from starting material to optimized compound [55]. This was in good agreement with increasing hydrogen bond acceptors favoring binding to GPCRs.

Model AAAR.2 could distinguish agonists from antagonists. Such mechanistic distinction is challenging, as A_2a_R agonists and antagonists often shared the same bicyclic adenine core [56]. Agonists and antagonists even engaged the same set of residues, such as Phe168, Ile274 and Asn253 as revealed by crystallographic studies [57-59]. The ribose ring structure is the key feature that differentiates agonists from antagonists [56]. As revealed by the co-crystal structures of A_2a_R and its agonist UK-432097, the ribose moiety was buried deeply into the binding pocket. The indole from a conserved Trp246 residue moved by ~1.9 Å to avoid clashing into the ribose ring. Such movement not only allowed additional contacts to be made with the ribose ring of the agonist, but also caused global movements to render the receptor’s transition into active form. Intriguingly, model AAAR.2 focused primarily on the adenine moiety (with exception of 1 hydrogen acceptor), hence might limit the identification of antagonists from agonists. Yet AAAR.2 still yielded above 70% sensitivity and specificity against a collection of 75 known A_2a_R agonists and antagonists. It is important to note that both sensitivity and specificity may be under-estimated. 16 out of the 46 antagonists were weak against A_2a_R, i.e., pIC_50_ < 5.3. These antagonists may be missed, within standard error, when the cutoff value for pIC_50_ was set to be 5.0. Such “omissions” may induce an underestimation of sensitivity, which could have been higher had the pIC_50_ values for all the antagonists were above 6.0. The specificity might also be higher than 70%. In our study, agonists identified as active by AAAR.2 were deemed as false positives, resulting in a false positive rate of ~30%. However, this might be too restrictive. Many agonists could also have antagonistic activities, as later demonstrated with the *in vitro* assay results for adenosine and regadenosine. Therefore the greater than 70% sensitivity and specificity were encouraging for this prospective application.

### The promise of utilization and interpretation of pharmacophore-sbased 3D QSAR in Safety

With large and diverse compound sets to generate various training sets and models, followed by thorough evaluation with a structurally different external set, 3D QSAR modeling could be used in safety, either as a pre-screen or to support detailed structural activity analysis against liability targets. To this end, it is important to measure the chemical similarities between query compounds and training/test compounds. Chemical similarity analysis is not simply to determine whether the query compound is suitable for the model or not. Rather it will help guide result interpretation, especially in safety screening.

This concept is best illustrated in **Fig 7**, which was divided into 4 areas including true positives, true negatives, false positives, and false negatives based on the results of *in vitro* and *in silico* assays against the subset of 75 A_2a_R ligands. The 5 compounds that are the most similar to training/test compounds (similarities ≥ 0.22) all fell into the section of true positives (**Figs 7A** and **7B**). For the 11 compounds whose similarity index ranged from 0.14 to 0.22 to training/test compounds, false positives and negatives began to appear, i.e., 2 were false positives and 2 were false negatives (**Fig 7C**). When similarity index dropped below 0.14, false positives and false negatives increased (**Fig 7D**). Encouragingly, 7 compounds (xanithinol, PSB603, PSB36, MRS1065, MRS1084, tonapofylline and sakuranetin), were also predicted positives despite their low similarity (similarity <0.1). The similarity was obtained from binary fingerprints, with no consideration for 3-dimensional or the pharmacophore features of the compounds. The success in prediction of these 7 compounds highlighted the advantage of 3D pharmacophore modeling over the 2D chemical features.

**Fig 7.**
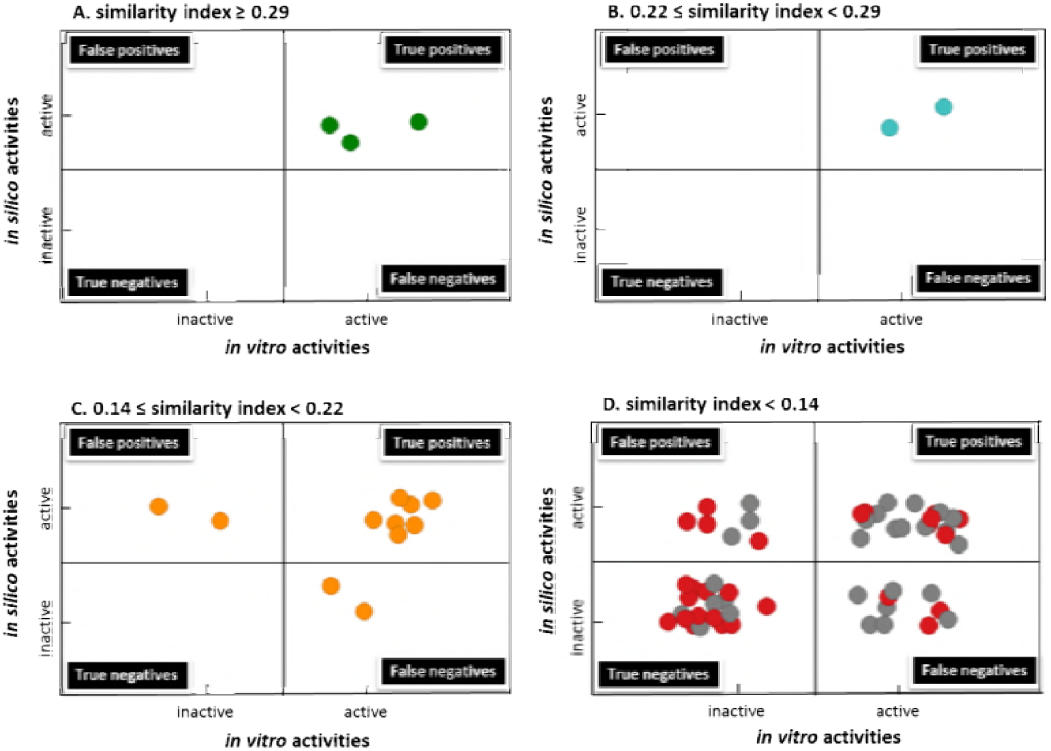
The performance of pharmacophore-based 3D QSAR modeling results in comparison to *in vitro* activities, when the similarities of the binary fingerprint between the query compound and the training/test compounds are ≥ 0.29 (**A**), between 0.22 to 0.29 (**B**), between 0.14 and 0.22 (**C**), and < 0.14 (**D**). In **D**, red dots indicated that similarity ranges between 0.10 to 0.14; grey dots indicated that similarity was below 0.10.

It is also important to note that that true negatives appeared with primarily low similarity compounds. This is in agreement with the hypothesis that chemicals with similar features may share similar targets. As such QSAR model could generally predict negatives with higher confidence, which is especially valuable in safety profiling versus efficacy profiling. If molecules with undesirable properties can be ruled out using virtual screening approaches, significant resources can be saved where only “prescreened” molecules are advanced to more costly *in vitro* screens. Among the 27 compounds whose similarity are under 0.1 in comparison to training/test compounds, 7 were true positives and 13 were true negatives. Among these low similarity compounds, 3 and 4 are false negatives and false positives respectively, giving 11% and 15% false negative and false positive rates. In safety, false positives could be later overruled with negative results obtained from the follow up *in vitro* assays. However, false negatives are more problematic. The 3 false negatives were MRS1191, FK453, and LUF5981. MRS1191 and FK453 were weak antagonists, with pIC_50_ values of 5.0 and 5.8, respectively. LUF5981, despite being a relatively potent antagonist with pIC_50_ values of 6.7, was reported to occupy the A_2a_R binding pocket in a different fashion [60,61]. Such ligand(s) with shifted binding position displayed the limitation of pharmacophore-based 3D QSAR model. Nonetheless, the false positive and negative rates were still within a tolerable range even for the standard of *in vitro* assays.

Therefore, when deploying a pharmacophore model, although comparing chemical similarity is important, one should not be discouraged from using the model simply because of low similarity to training/test compounds, particularly due to the nature and emphasis of a safety (pre)screening. The model would still be valuable when similarities are low, as demonstrated from compounds that were least similar to training and test sets. More importantly, chemical similarity helps guide the interpretation of predicted data. For the utilization of a virtual screen in safety, positive prediction outcomes could be interpreted with confidence when similarity is high (in our case, when similarity > 0.22). False positives and negatives should be expected as they coincide with decreasing similarity (e.g., from 0.22 to 0.14). When similarity is very low (e.g. < 0.1), negative predictions may be interpreted with confidence based on the principal “chemical with similar structures may bind to similar targets”. The similarity cut-off values should be established with carefully curated external set prior to the prospective utilization. In our case, we used the subset of 75 A_2a_R ligands. *In vitro* assay follow up is highly recommended in the following 2 cases. One is when the similarity is low yet positives are predicted, the other is when the similarity is high yet negatives are predicted.

In summary, we presented a study to evaluate the possibility of incorporating *in silico* screening in the arena of safety. Instead of being carried out as a retrospective exercise, we focused on prospective utilization in safety screening. Our study was designed with several distinct features, such as generating multiple models from various training and test sets, and utilization of structurally different external set as well as a larger and more diverse set of compounds from the real world. When integrating pharmacophore-based 3D QSAR in safety, we recommend the following based on our analysis. First, large and diverse compound set should be used to generate the model. Addition of extra compounds and data from publication and public database will help enrich the diversity of training and test sets, hence increase the prospects for future utilization of the model in broadened chemical space. Second, multiple training and test sets should be generated, and accordingly multiple models (possibly containing different number of pharmacophore sites) should be evaluated. Third, thorough evaluation using a structurally different external set with multiple models is important to evaluate the performance against new chemotypes. The external set also helps establish the similarity cutoff values for future prospective utilization of the model. Last, the interpretation of prediction outcome should be viewed in combination with similarity analysis of query compound(s) and training compounds, which will also help to prioritize the subsequent *in vitro* follow-ups. With these steps this detailed case study demonstrated that an otherwise limited ligand-based QSAR approach may be nicely integrated into the *in vitro* safety profiling, either as a pre-screen prior to *in vitro* assays (for new chemotypes before they are even made) or to support detailed SAR against liability targets. Neither is aimed at the discovery of new chemical series, rather, the value of pharmacophore-based 3D QSAR model lies in helping to “design away” from liability targets during Drug Development

### Supporting Information

An enlarged version of Fig 2, as well as an example of QSAR visualization of positive and negative regression coefficients for active vs inactive molecules.

## Author Contributions

The manuscript was written through contributions of all authors. / All authors have given approval to the final version of the manuscript.

## ACKNOWLEDGMENT

The authors thank Dr. Hua Gao (Molecular Engineering, Amgen) for his suggestions on exploring the similarities between query compound and training/test sets. The authors also thank Dr. Wei Zhang and Dr. Mark R. Fielden (CBSS, Amgen) for their insightful reviews on this manuscript.

## ABBREVIATIONS

A_2a_R, adenosine receptor 2a; QSAR, quantitative structure activity relationship; HTS, high-throughput screen

PLS, partial least square; RMSE, Root Mean Square Error ; SD, standard deviation of regression; MOA, mode of action

## REFERENCES

1. Bowes J, Brown AJ, Hamon J, Jarolimek W, Sridhar A, et al. (2012) Reducing safety-related drug attrition: the use of in vitro pharmacological profiling. Nat Rev Drug Discov 11: 909–922.

2. Hamon J, Whitebread S, Techer-Etienne V, Le Coq H, Azzaoui K, et al. (2009) In vitro safety pharmacology profiling: what else beyond hERG? Future Med Chem 1: 645–665.

3. Redfern WS, Wakefield ID, Prior H, Pollard CE, Hammond TG, et al. (2002) Safety pharmacology--a progressive approach. Fundam Clin Pharmacol 16: 161–173.

4. Whitebread S, Dumotier B, Armstrong D, Fekete A, Chen S, et al. (2016) Secondary pharmacology: screening and interpretation of off-target activities - focus on translation. Drug Discov Today 21: 1232–1242.

5. Whitebread S, Hamon J, Bojanic D, Urban L (2005) Keynote review: in vitro safety pharmacology profiling: an essential tool for successful drug development. Drug Discov Today 10: 1421–1433.

6. Concil NR (2007) Toxicity testing in the 21st century: a vision and a strategy. Washington D. C.: The National Academies Press.

7. Warren I, Eftim S (2013) Use of Publicly Avaiable Data in Risk Assessment. In: Fowler BA, editor. Computational Toxicology: Methods and Applications for Risk Assessment. Waltham, MA: Academic Press. pp. 151–167.

8. Gibb S (2008) Toxicity testing in the 21st century: a vision and a strategy. Reprod Toxicol 25: 136–138.

9. Urban L, Whitebread S, Hamon J, Mikhailov D, Azzaoui K (2012) Screening for Safety Related Off-Target Activities. In: Peters J, editor. Polypharmacology in Drug Discovery. Hoboken, NJ: John Wiley & Sons, INC. pp. 15–46.

10. Mansouri K, Judson RS (2016) In Silico Study of In Vitro GPCR Assays by QSAR Modeling. Methods Mol Biol 1425: 361–381.

11. Muster W, Breidenbach A, Fischer H, Kirchner S, Muller L, et al. (2008) Computational toxicology in drug development. Drug Discov Today 13: 303–310.

12. Vedani A, Smiesko M (2009) In silico toxicology in drug discovery - concepts based on three-dimensional models. Altern Lab Anim 37: 477–496.

13. Vedani A, Dobler M, Lill MA (2006) The challenge of predicting drug toxicity in silico. Basic Clin Pharmacol Toxicol 99: 195–208.

14. Sprous DG, Palmer RK, Swanson JT, Lawless M (2010) QSAR in the pharmaceutical research setting: QSAR models for broad, large problems. Curr Top Med Chem 10: 619–637.

15. Todeschini R, Consonni V (2008) Handbook of Molecular Descriptors; Mannhold R, Kubinyi H, Timmerman H, editors. Weinheim, Germany

16. Cherkasov A, Muratov EN, Fourches D, Varnek A, Baskin, II, et al. (2014) QSAR modeling: where have you been? Where are you going to? J Med Chem 57: 4977–5010.

17. Fourches D, Muratov E, Tropsha A (2010) Trust, but verify: on the importance of chemical structure curation in cheminformatics and QSAR modeling research. J Chem Inf Model 50: 1189–1204.

18. Williams AJ, Ekins S (2011) A quality alert and call for improved curation of public chemistry databases. Drug Discov Today 16: 747–750.

19. Cronin MTD, Schultz TW (2003) Pitfalls in QSAR. Journal of Molecular Structure: THEOCHEM 622: 39–51.

20. Nigsch F, Bender A, Jenkins JL, Mitchell JB (2008) Ligand-target prediction using Winnow and naive Bayesian algorithms and the implications of overall performance statistics. J Chem Inf Model 48: 2313–2325.

21. Raies AB, Bajic VB (2016) In silico toxicology: computational methods for the prediction of chemical toxicity. Wiley Interdiscip Rev Comput Mol Sci 6: 147–172.

22. The OECD QSAR toolbox.

23. Keiser MJ, Roth BL, Armbruster BN, Ernsberger P, Irwin JJ, et al. (2007) Relating protein pharmacology by ligand chemistry. Nat Biotechnol 25: 197–206.

24. Patlewicz G, Jeliazkova N, Gallegos Saliner A, Worth AP (2008) Toxmatch-a new software tool to aid in the development and evaluation of chemically similar groups. SAR QSAR Environ Res 19: 397–412.

25. Patlewicz G, Jeliazkova N, Safford RJ, Worth AP, Aleksiev B (2008) An evaluation of the implementation of the Cramer classification scheme in the Toxtree software. SAR QSAR Environ Res 19: 495–524.

26. Richard AM, Yang C, Judson RS (2008) Toxicity data informatics: supporting a new paradigm for toxicity prediction. Toxicol Mech Methods 18: 103–118.

27. Valerio LG, Jr. (2009) In silico toxicology for the pharmaceutical sciences. Toxicol Appl Pharmacol 241: 356–370.

28. Fredholm BB, Ap IJ, Jacobson KA, Klotz KN, Linden J (2001) International Union of Pharmacology. XXV. Nomenclature and classification of adenosine receptors. Pharmacol Rev 53: 527–552.

29. Fredholm BB, Cunha RA, Svenningsson P (2003) Pharmacology of adenosine A2A receptors and therapeutic applications. Curr Top Med Chem 3: 413–426.

30. Dai SS, Zhou YG, Li W, An JH, Li P, et al. (2010) Local glutamate level dictates adenosine A2A receptor regulation of neuroinflammation and traumatic brain injury. J Neurosci 30: 5802–5810.

31. Dunwiddie TV, Masino SA (2001) The role and regulation of adenosine in the central nervous system. Annu Rev Neurosci 24: 31–55.

32. Fredholm BB, Chen JF, Cunha RA, Svenningsson P, Vaugeois JM (2005) Adenosine and brain function. Int Rev Neurobiol 63: 191–270.

33. Fredholm BB, Chen JF, Masino SA, Vaugeois JM (2005) Actions of adenosine at its receptors in the CNS: insights from knockouts and drugs. Annu Rev Pharmacol Toxicol 45: 385–412.

34. Shen HY, Chen JF (2009) Adenosine A(2A) receptors in psychopharmacology: modulators of behavior, mood and cognition. Curr Neuropharmacol 7: 195–206.

35. Huang ZL, Urade Y, Hayaishi O (2011) The role of adenosine in the regulation of sleep. Curr Top Med Chem 11: 1047–1057.

36. Eltzschig HK, Sitkovsky MV, Robson SC (2012) Purinergic signaling during inflammation. N Engl J Med 367: 2322–2333.

37. Chen J, Eltzschig HK, Fredholm BB (2013) Adenosine receptors as drug targets — what are the challenges? Nat Rev Drug Discov 12: 265–286.

38. Gessi S, Merighi S, Sacchetto V, Simioni C, Borea PA (2011) Adenosine receptors and cancer. Biochim Biophys Acta 1808: 1400–1412.

39. Mustyala KK, Chitturi AR, Naikal James PS, Vuruputuri U (2012) Pharmacophore mapping and in silico screening to identify new potent leads for A(2A) adenosine receptor as antagonists. J Recept Signal Transduct Res 32: 102–113.

40. Scior T, Medina-Franco JL, Do QT, Martinez-Mayorga K, Yunes Rojas JA, et al. (2009) How to recognize and workaround pitfalls in QSAR studies: a critical review. Curr Med Chem 16: 4297–4313.

41. Wishart DS, Knox C, Guo AC, Shrivastava S, Hassanali M, et al. (2006) DrugBank: a comprehensive resource for in silico drug discovery and exploration. Nucleic Acids Res 34: D668–672.

42. Bento AP, Gaulton A, Hersey A, Bellis LJ, Chambers J, et al. (2014) The ChEMBL bioactivity database: an update. Nucleic Acids Res 42: D1083–1090.

43. Alexander SP, Kelly E, Marrion N, Peters JA, Benson HE, et al. (2015) The Concise Guide to PHARMACOLOGY 2015/16: Overview. Br J Pharmacol 172: 5729–5743.

44. Sastry M, Lowrie JF, Dixon SL, Sherman W (2010) Large-scale systematic analysis of 2D fingerprint methods and parameters to improve virtual screening enrichments. J Chem Inf Model 50: 771–784.

45. Greenwood JR, Calkins D, Sullivan AP, Shelley JC (2010) Towards the comprehensive, rapid, and accurate prediction of the favorable tautomeric states of drug-like molecules in aqueous solution. J Comput Aided Mol Des 24: 591–604.

46. Shelley JC, Cholleti A, Frye LL, Greenwood JR, Timlin MR, et al. (2007) Epik: a software program for pK(a) prediction and protonation state generation for drug-like molecules. J Comput Aided Mol Des 21: 681–691.

47. Harder E, Damm W, Maple J, Wu C, Reboul M, et al. (2016) OPLS3: A Force Field Providing Broad Coverage of Drug-like Small Molecules and Proteins. J Chem Theory Comput 12: 281–296.

48. Dixon SL, Smondyrev AM, Knoll EH, Rao SN, Shaw DE, et al. (2006) PHASE: a new engine for pharmacophore perception, 3D QSAR model development, and 3D database screening: 1. Methodology and preliminary results. J Comput Aided Mol Des 20: 647–671.

49. Polanski J, Bak A, Gieleciak R, Magdziarz T (2006) Modeling robust QSAR. J Chem Inf Model 46: 2310–2318.

50. Fan F, Hu R, Munzli A, Chen Y, Dunn RT, 2nd, et al. (2015) Utilization of human nuclear receptors as an early counter screen for off-target activity: a case study with a compendium of 615 known drugs. Toxicol Sci 145: 283–295.

51. Dixon SL, Smondyrev AM, Rao SN (2006) PHASE: a novel approach to pharmacophore modeling and 3D database searching. Chem Biol Drug Des 67: 370–372.

52. Dixit A, Verkhivker GM (2011) The energy landscape analysis of cancer mutations in protein kinases. PLoS One 6: e26071.

53. Gregori-Puigjane E, Mestres J (2008) A ligand-based approach to mining the chemogenomic space of drugs. Comb Chem High Throughput Screen 11: 669–676.

54. Katritch V, Cherezov V, Stevens RC (2012) Diversity and modularity of G protein-coupled receptor structures. Trends Pharmacol Sci 33: 17–27.

55. Morphy R (2006) The influence of target family and functional activity on the physicochemical properties of pre-clinical compounds. J Med Chem 49: 2969–2978.

56. Xu F, Wu H, Katritch V, Han GW, Jacobson KA, et al. (2011) Structure of an agonist-bound human A2A adenosine receptor. Science 332: 322–327.

57. Jaakola VP, Ijzerman AP (2010) The crystallographic structure of the human adenosine A2A receptor in a high-affinity antagonist-bound state: implications for GPCR drug screening and design. Curr Opin Struct Biol 20: 401–414.

58. Jaakola VP, Lane JR, Lin JY, Katritch V, Ijzerman AP, et al. (2010) Ligand binding and subtype selectivity of the human A(2A) adenosine receptor: identification and characterization of essential amino acid residues. J Biol Chem 285: 13032–13044.

59. Kim J, Wess J, van Rhee AM, Schoneberg T, Jacobson KA (1995) Site-directed mutagenesis identifies residues involved in ligand recognition in the human A2a adenosine receptor. J Biol Chem 270: 13987–13997.

60. Chang LC, von Frijtag Drabbe Kunzel JK, Mulder-Krieger T, Westerhout J, Spangenberg T, et al. (2007) 2,6,8-trisubstituted 1-deazapurines as adenosine receptor antagonists. J Med Chem 50: 828–834.

61. Yang X, Dong G, Michiels TJ, Lenselink EB, Heitman L, et al. (2016) A covalent antagonist for the human adenosine A2A receptor. Purinergic Signal.

